# Analysis of proteome adaptation reveals a key role of the bacterial envelope in starvation survival

**DOI:** 10.1101/2022.05.18.492425

**Authors:** Severin Schink, Constantin Ammar, Yu-Fang Chang, Ralf Zimmer, Markus Basan

**Affiliations:** Systems Biology Department, Harvard Medical School, 200 Longwood Ave, 02115 MA, USA; Institute of Informatics, Ludwig-Maximilians-Universität München, Amalienstr. 17, 80333, Munich, Germany

## Abstract

Bacteria reorganize their physiology upon entry to stationary phase. What part of this reorganization improves starvation survival is a difficult question, because the change in physiology includes a global reorganization of the proteome, envelope and metabolism of the cell. In this work, we used several trade-offs between fast growth and long survival to statistically score over 2000 *E. coli* proteins for their global correlation with death rate. The combined ranking allowed us to narrow down the set of proteins that positively correlate with survival and validate the causal role of a subset of proteins. Remarkably, we found that important survival genes are related to the cell envelope, i.e., periplasm and outer membrane, because maintenance of envelope integrity of *E. coli* plays a crucial role during starvation. Our results uncover a new protective feature of the outer membrane that adds to the growing evidence that the outer membrane is not only a barrier that prevents abiotic substances from reaching the cytoplasm, but essential for bacterial proliferation and survival.

**Standfirst text:** A trade-off between the two major modes of bacterial lifestyle, growth and starvation can be explained by bacteria investing resources into the cell envelope to make it impermeable to ions, which improves their lifespan but comes at the expense of slowing down growth.

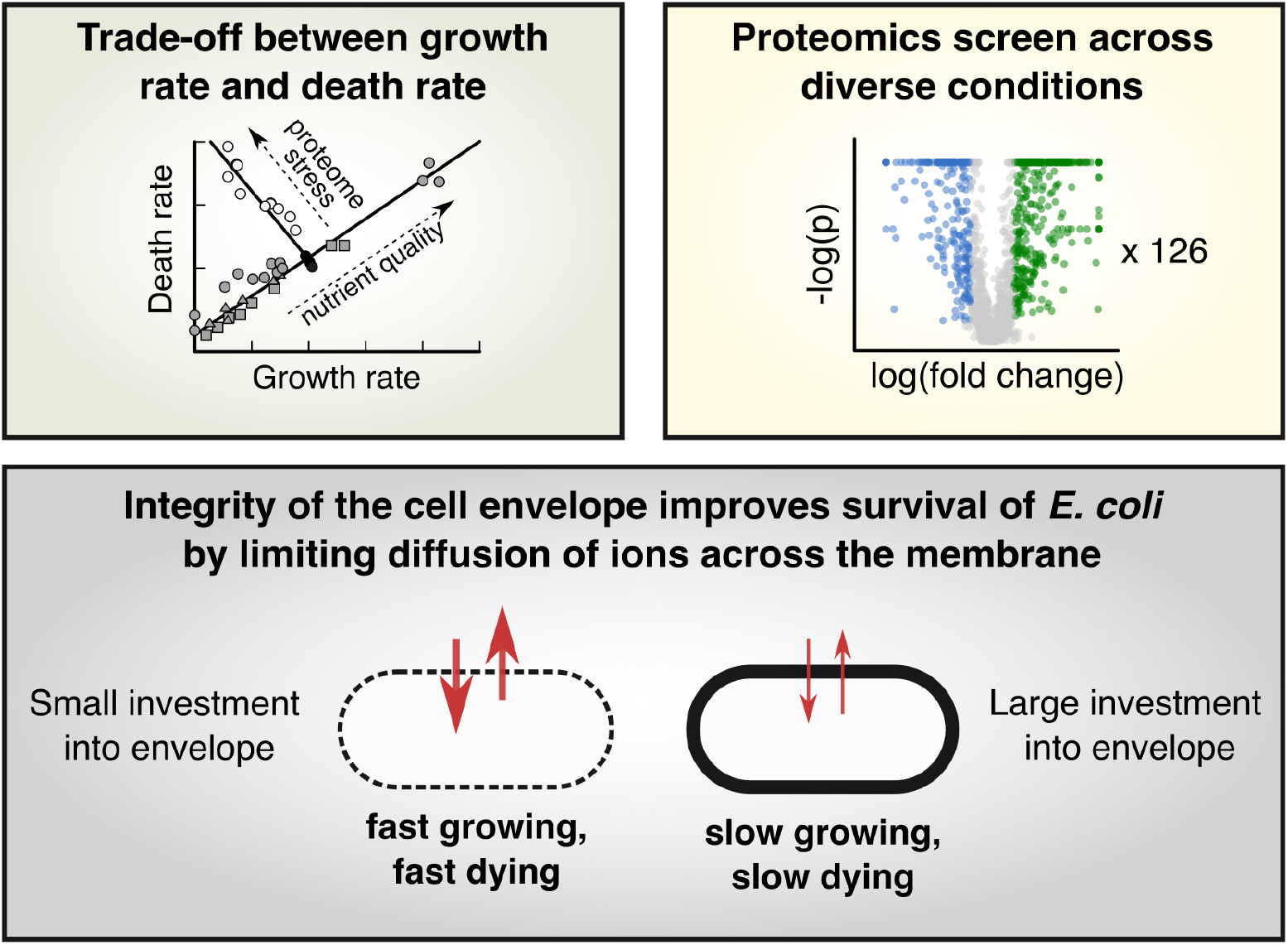

**Highlights:**

- A trade-off between growth rate and death rate confines fitness of bacteria across environments.
- Analysis of proteome signatures in 126 conditions across five independent perturbations reveals the cell envelope as a key determinant of death rate.
- The trade-off can be abolished by changing environment to a low-salt, but osmo-balanced medium where cell envelope integrity is not limiting.

## Introduction

Nutrient limitation is a defining part of the lifecycle of microorganisms. In the absence of external nutrients, the only energy source for heterotrophic organisms is either internal storage (Fung and Kwong, 2013; Hengge-Aronis and Fischer, 1992; Strange, 1968) or nutrients retrieved by recycling dead biomass (Steinhaus and Birkeland, 1939; Zambrano and Kolter, 1996; Schink et al., 2019; Finkel, 2006). These finite energy sources can be temporarily used to maintain the cell, but will eventually deplete, exposing cells to a slow deterioration process driven by entropic forces. It was previously shown that the survival kinetics during starvation, i.e. how many cells will be still alive after a certain time, is determined by the consumption rate of these nutrients, called the maintenance rate, and the availability of nutrients (Schink et al., 2019). Because maintenance rate is a major determinant of bacterial lifespan, we expect bacteria to minimize their maintenance cost to maximize fitness in starvation.

But what determines how much maintenance bacteria need to perform, and how they can reduce it is still largely unclear. The question of maintenance is particularly puzzling, because sporulating organisms can minimize the number of active processes without losing their viability. The maintenance rate of non-sporulating bacteria is similarly not an engraved biophysical constant. Instead, *Escherichia coli*, for example, can decrease its maintenance rate and death rate over 5-fold in response to environmental cues sensed during the prior growth phase (Biselli et al., 2020).

But identifying the part of the adaptation that determines lifespan is tricky as the bacteria globally reorganize their proteome (Hui et al., 2015; Schmidt et al., 2016), macromolecular composition (Bremer and Dennis, 2008; Schaechter et al., 1958) and even morphology (Schaechter et al., 1958), when nutrients get depleted and growth slows down. This means that most properties of bacteria correlate or anti-correlate with lifespan. In addition, while several regulators and genes have been identified that react to nutrient depletion and are essential for survival, such as the general stress response regulator *rpoS* (Hengge-Aronis and Fischer, 1992), cAMP-Crp (Makman and Sutherland, 1965; Notley-McRobb et al., 1997), stringent response (Magnusson et al., 2003; Irving et al., 2021) or DNA protection in starvation *dps* (Almirón et al., 1992), there is no evidence that these systems are the limiting factors for survival.

In this paper, we analyzed diverse growth conditions and find that they heavily influence lifespan after nutrients run out. Due to the diversity in proteomic responses, we can use this information to narrow down the set of proteins that correlate with better survival and validate which parts of the adaptation determine lifespan of *E. coli* in starvation.

## Results

### Proteome analysis identifies proteins and processes that correlate with long lifespan

We studied *Escherichia coli* K-12, first grown in single carbon source N^+^C^+^ minimal medium, followed by a wash and resuspension in medium with the carbon source missing. The washing step introduces bacteria suddenly to starvation and prevents their complete adaptation. As a result, during the ensuing starvation, viability decreases exponentially (Phaiboun et al., 2015; Schink et al., 2019), with death rate depending on the previous growth condition. *E. coli* grown on carbon substrates supporting faster growth die faster than those on poor carbon substrates (Biselli et al., 2020), see Fig. 1A (‘blue’). Alternatively, letting *E. coli* adapt to stationary phase on glucose minimal medium results in a death rate similar to sudden starvation from very low growth rates (Fig. 1A, open symbols placed at growth rate zero). We suspected that changes in the proteome composition, which is known to vary highly across growth conditions (Hui et al., 2015; Schmidt et al., 2016), are at least partially responsible for the observed changes in death rate. However, in a single growth perturbation, such as varying the carbon substrates, over half of all proteins were up- or downregulated (Hui et al., 2015; Schmidt et al., 2016), making it impossible to pinpoint individual proteins or processes as being responsible.

**Figure 1.**
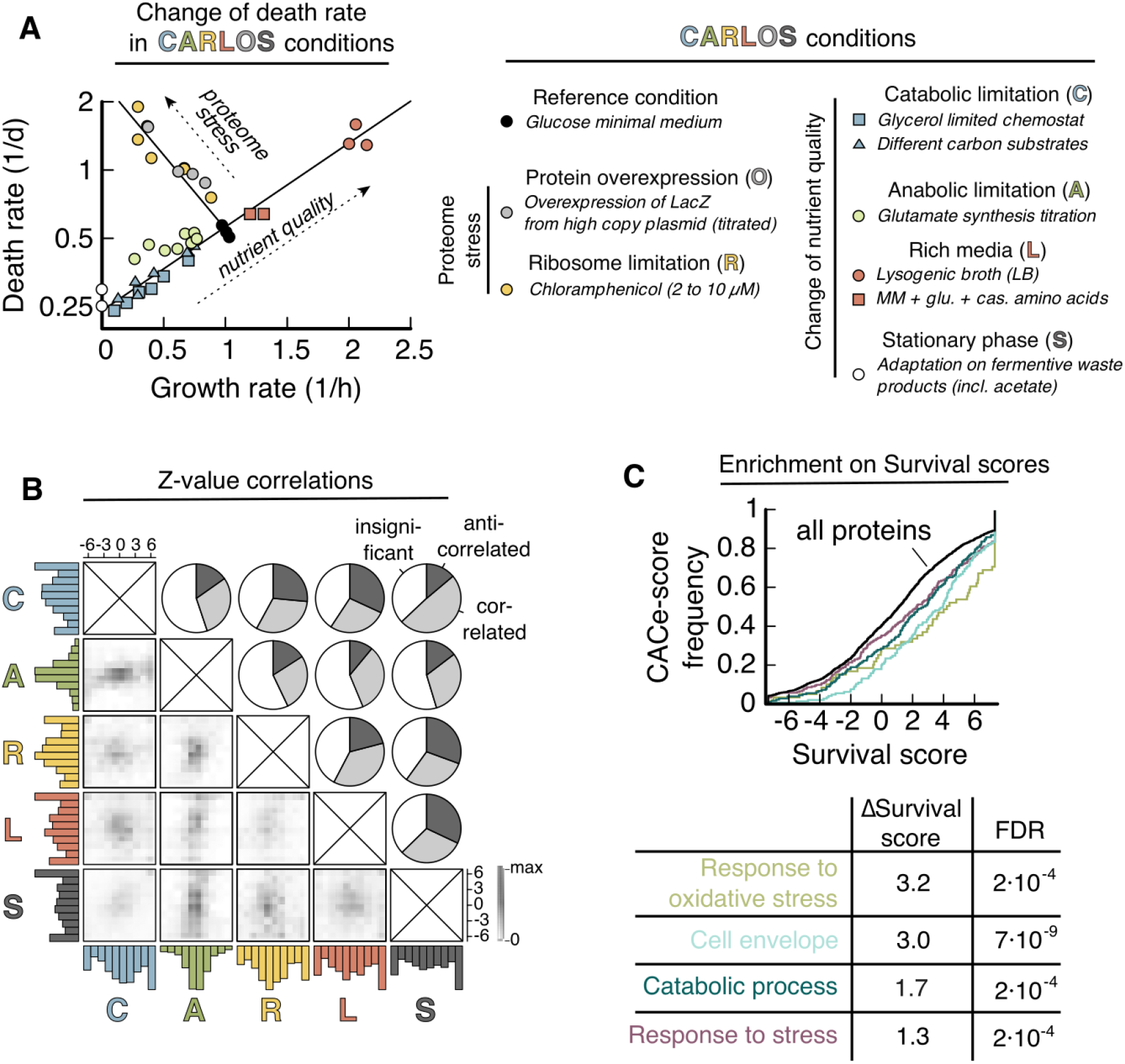
Proteome composition and death rates in CARLOS conditions. (A) Death rate versus pre-starvation growth rate on a semilog-plot. CARLOS conditions are defined in legend. CARLOS conditions that perturb nutrient quality (C, A, L and S) lead to an exponential increase of death rate. Conditions that stress the proteome (R and O) show an orthogonal exponential increase with slower growth. Each data point is a single experiment. Chemostat (C) and parts of (L) are from Ref. (Biselli et al., 2020). See Table S1 for data. (B) Pair-wise comparison of protein Z-values between conditions. Histograms of Z-values of individual conditions (colored) and grey values of the 2D histogram between pairs of conditions are normalized to their respective maximum value. The fraction of proteins that correlate, anti-correlate or are insignificant is calculated by thresholding in the 2D histogram, see Fig. EV1A for an illustration. (C) Z-values are merged into a single ‘Survival score’ for each protein that measures correlation of abundance with survival across all experiments. A detailed description of the construction of the score is given in Box 1. Several GO-processes are significantly enriched (FDR levels of the Benjamini-Hochberg step-up procedure) towards higher scores. ΔSurvival score is calculated as the difference in median Survival score between a process and the background of all proteins.

To narrow down the search for the proteins that determine lifespan, we decided to use additional growth conditions that show different changes in proteome. We identified a total of six independent perturbations, which we collectively call ‘CARLOS conditions’ that affect proteome composition during growth and death rate after washing and resuspension in carbon-free medium. Among these, those that reflect a change in nutrient availability (C: <u>c</u>atabolic limitation, A: <u>a</u>nabolic limitation, L: rich media (<u>L</u>B) and S: <u>s</u>tationary phase) collapse onto a single exponential increase of death rate with growth rate, while conditions that perturb the proteome by expression of excess proteins (R: ribosome limitation, O: LacZ <u>o</u>verexpression) fall onto an orthogonal exponential increase (Fig. 1A).

CARLOS conditions result in substantial changes in proteome composition. We analyzed these changes in the proteome using published proteomics data from three different repositories (Data ref: Houser et al., 2015; Data ref: Hui et al., 2015; Data ref: Schmidt et al., 2016), which individually cover a subset of the CARLOS conditions (C, L & S in (Data ref: Schmidt et al., 2016), C, A & R in (Data ref: Hui et al., 2015) and S in (Data ref: Houser et al., 2015)). To be able to compare proteins across conditions and across data sets, we created a composite statistical score, termed “Survival Score”, explained in detail in Box 1 and the Methods section.

**Box 1.**
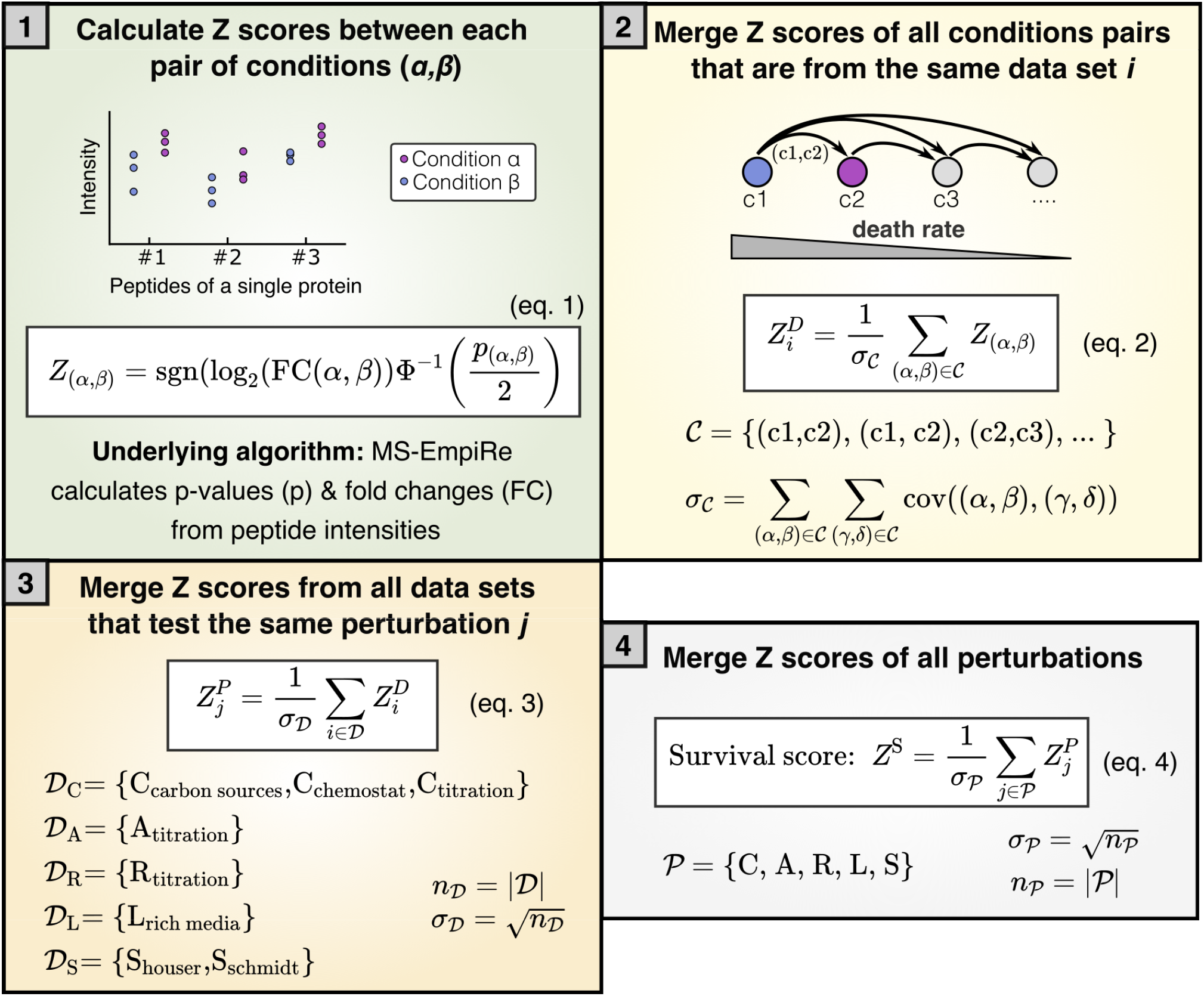
Calculation of Survival scores from multi-condition proteomics data sets. The survival scores *Z*^*s*^ quantifies how well a protein correlates with lifespan over all CARLS perturbations and is calculated in the steps listed below. Jupyter Notebooks implementing the procedure below are provided in the SI (see also data availability section). 1) Using the MS-Empire statistical software package, we calculated p-values and fold-changes (FC) of proteins between a pair of conditions (*α, β*) based on the underlying peptide intensities across replicates (blue, purple). We use this information to calculate the Z score of a pair of conditions, *Z*_(*α,β*)_, in eq. (1) using the direction of the fold-change, where ‘sgn’ is the sign function, and the significance of the regulation, using the inverse cumulative standard normal distribution function Φ^−1^. 2) Next, we merge all *Z*_(*α,β*)_ that are from the same data set *𝒞*, into a new Z value of a data set 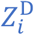 in eq. (2), where the subscript *i* stands for the specific data set. An example of a data set are *Z*_(*α,β*)_ scores of different carbon sources derived from the same data repository. The pairs of conditions (*α, β*) are sorted from high to low death rate, such that a positive Z-value corresponds to correlation with lower death rate. Because the merged score 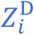 should follow a standard normal distribution with standard deviation of 1, merging *Z*_(*α,β*)_ scores requires normalization by the standard deviation across the whole data set, *σ*_*𝒞*_. The standard deviation can be derived from the covariance matrix, where cov/(*α, β*), (*γ*, δ)4 is 1 if (*α* = *γ* AND *β* = *δ*), 0.5 if (*α* = *γ* XOR *β* = *δ*) and 0 otherwise (Ammar et al., 2019). 3) Next, we merge all 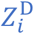 that belong to the same perturbation into a single score for each perturbation 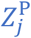, where the subscript *j* stands for any of the CARLS conditions. Because data sets are independent of each other, the standard deviation *σ*_𝒟_ depends only on the number of data sets *n*_𝒟_. For A, R and L there is only a single data set, such that 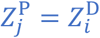. Fig. EV1 visualizes this merging step for C and S conditions. *4)* Finally, all 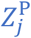 are merged into a single *Z*^S^, which we call ‘Survival score’. As in panel 3, the standard deviation *σ*_*𝒫*_ depends only on the number of perturbations *n*_*𝒫*_ that are measured.

In short, we compare protein abundance levels between data sets and assign a positive score (in the form of a Z-value), if the protein is correlated with better survival and a negative score if the protein is anticorrelated (Box 1, step 1). We collect these scores over the variety of conditions and data sets and iteratively merge the scores, while correcting for statistical dependencies and ignoring missing values (steps 2-4).

Z-values calculated from individual data sets of similar perturbations or different sources (Box 1, step 2) are highly correlated (Fig. EV1A&B) and are used to create merged Z-values for each protein (Fig. EV1C & Box 1, step 3). Histograms of these merged Z-values on the condition level (step 3) are shown as colored 1D histograms in Fig. 1B, normalized to the highest value. We had to exclude ‘O, LacZ overexpression’ from this analysis due to lack of sufficient statistical data, shortening CARLOS to CARLS, but we will integrate qualitative information from this condition below.

Next, we compared Z-values of the conditions in a pair-wise manner by plotting 2D histograms of Z-values detected in both respective conditions (Fig. 1B, bottom left of matrix). By classifying proteins that score above |*Z*| > 1.28, which corresponds to *p* < 0.1, as significantly correlated or anti-correlated, depending on the direction, we confirmed that proteome changes across different conditions are partially orthogonal to each other (Fig. 1B, top right of matrix). This means that each condition has additional information that can help us narrow down the target set of proteins. Finally, we merged Z-values of the individual CARLS conditions into a single ‘Survival score’ that measures the global correlation strength of proteins with lifespan (Box 1 step 4). The resulting list of CARLS Z-values and ‘Survival scores’ for over 2000 proteins is available in Table S2.

Fractions of the proteome determining death rate can be either beneficial or harmful. To understand which of these fractions is dominant, we used the ‘O’ condition, where overexpression of a useless protein leads to a decrease in the abundance of virtually all proteins simultaneously (Fig. EV2). This condition led to a 5-fold increase in death rate (Fig. 1A, gray symbols), as we would expect for largely beneficial proteins determining death rate. Thus, we restricted our further analysis to positive Survival scores, where a higher abundance correlates with longer lifespan across conditions.

Finally, we used Survival scores of all proteins measured in at least 3 of the 5 CARLS conditions to identify gene ontology (GO) processes and cellular compartments that contain proteins with significantly larger Survival scores than average. Using a Kolmogorov-Smirnov test, we evaluated enrichment of proteins in GO-processes and compartments, which includes ‘oxidative stress response’, ‘response to stress’, ‘catabolic process’ and ‘cell envelope’ as some of the highest scoring GO-processes (Fig. 1C, see Table S3 for full list). Note that GO-processes are not mutually exclusive and can contain the same proteins, which is particularly true for ‘oxidative stress response’, which substantially overlaps with ‘response to stress’.

### Death rate depends on the integrity of the cell envelope

We next wanted to test if any of these processes/compartments causally affect bacterial lifespan. The response to oxidative stress had the highest Survival score in our analysis, with proteins such as catalases KatE (Survival score: 7.3) and KatG (7.3), superoxide dismutase sodA (6.5), and hydroperoxidase AhpC (6.2), all of which are essential to *E. coli’s* adaptation to stress (Farr et al., 1988; Farr and Kogoma, 1991; Dukan and Nyström, 1999; Chiang and Schellhorn, 2012), correlating well with survival across CARLOS conditions. Therefore, we first tested whether oxidative stress limits lifespan of starving bacteria by starving bacteria in anaerobic conditions, where oxidative damage should be absent. It was previously reported that starved *E. coli* suffer from protein carbonylation and that anaerobic conditions can prevent carbonylation and stop death (Dukan and Nyström, 1999). However, to our surprise, our results showed the opposite. We found that death in anaerobic conditions was even faster than in aerobic conditions (Fig. EV3A). This did not depend on whether growth prior to starvation was in aerobic or anaerobic conditions. Similarly, adding antioxidants glutathione, ascorbic acid or mercaptoethanol, which could prevent carbonylation during starvation, did not decrease the death rate of starvation *E. coli* (Fig. EV3B).

Also, inducing the ‘response to stress’ did not decrease death rate, despite the high score. Neither inducing its expression using heat shocks, low pH nor high osmolarity during growth, all of which significantly increase abundance of ‘response to stress’ proteins (Fig. EV4A-C), resulted in changes of death rate of *E. coli* in reference starvation conditions (Fig. EV4D). One possible explanation is that our wild-type strain NCM3722 is insensitive because one of the master regulators of the stress response, *rpoS*, contains a premature amber stop codon, which reduces expression of *rpoS* dependent genes (Mori et al., 2021). But restoring *rpoS* did not decrease death rate (Fig. EV4E). In contrast, knockouts of *rpoS* still showed significantly increased death rates compared to NCM3722 (Fig. EV4E), which indicates that the *rpoS* gene is sufficiently functional in our strain. Only a knock-out *rssB*, a gene responsible for degradation of RpoS (Muffler et al., 1996) lead to a significant decrease in death rate (Fig. EV4E), similar to previous reports (Fontaine et al., 2008). These conflicting observations warrant further investigation into physiological role of *rpoS* and *rssB* in starvation survival. However, it is known that *rpoS* knockouts do not remove the growth-death dependence in carbon limitation (Biselli et al., 2020) that forms the basis of our analysis.

Another high-scoring class of proteins identified by our analysis were proteins of the cell envelope, which includes the periplasm, cell wall and outer membrane (Fig. EV5 for graphical summary). Proteins associated with the cell envelope make up about 13% of the proteome in rich nutrient conditions, and up to 23% for bacteria adapted to stationary phase (Fig. 2A and Table S4). In addition to the GO analysis, we also noticed the cell envelope in a knock-out screen of high-scoring, non-essential genes. The fastest dying knock-out in the screen, Δ*lpp*, encoded a protein located in the cell envelope (Table S5). In contrast, knockouts of most other genes, especially those belonging to stress response and oxidative damage, had little or no impact of death rates in starvation (Table S5), further corroborating that many of them are non-limiting in our conditions.

**Figure 2.**
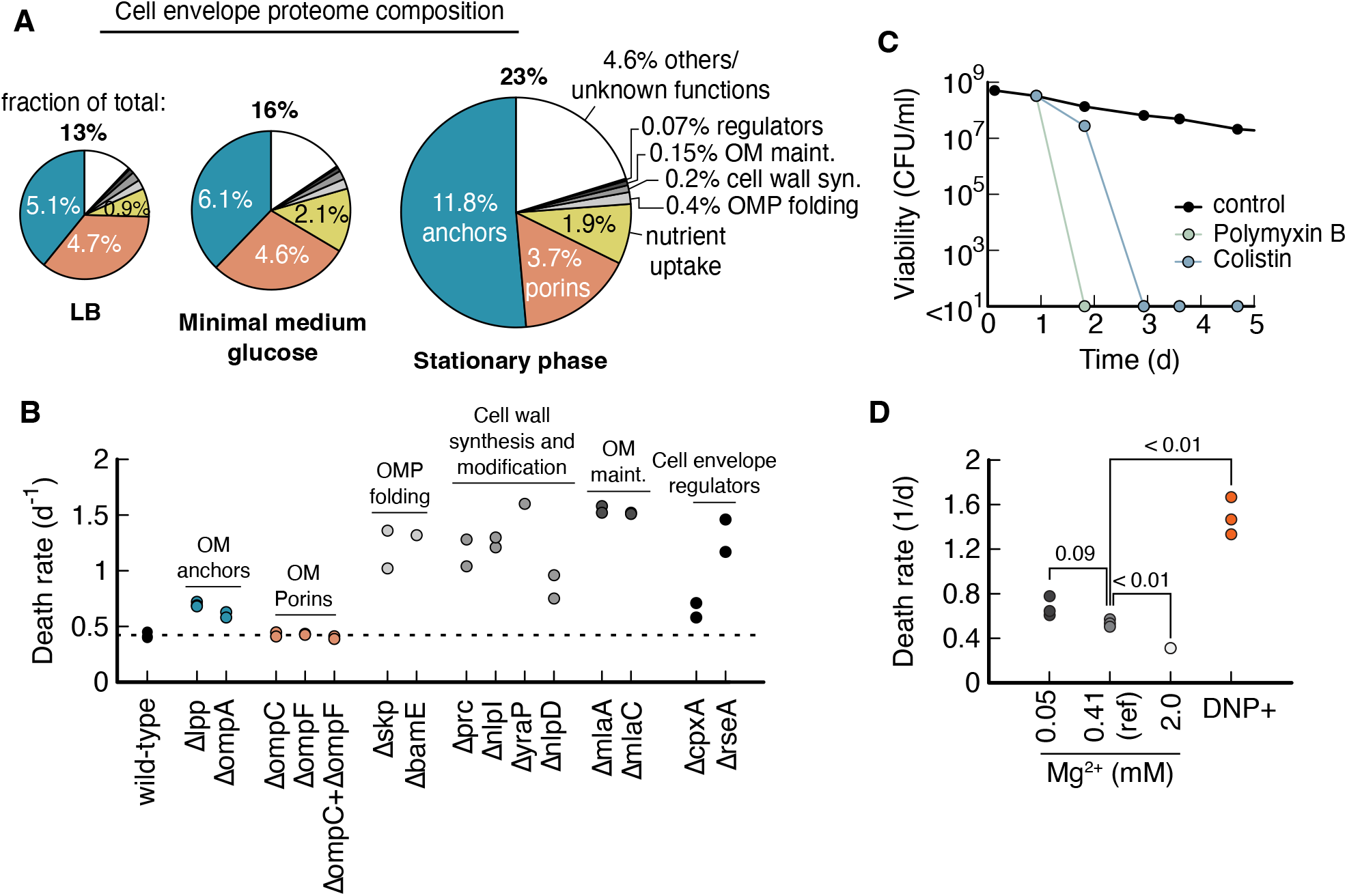
Cell envelope integrity plays a major role in starvation survival. (A) Proteome composition of the cell envelope (see Fig. EV5 for details on the cell envelope structure) for cultures during exponential growth on LB, glucose minimal medium and stationary phase. Size of pie indicates fraction of total proteome that is associated with envelope. Outer membrane anchors (Lpp and OmpA) constitute half of the cell envelope proteome. Major porins (OmpC and OmpF) and nutrient uptake (large number of binding proteins) constitute other major fractions. Quantification of abundancies are in Table S4 using data from Ref. (Data ref: Schmidt et al., 2016). (**B**) Knock-out of several genes with different functionalities in the cell envelope increase death rates. Growth in glycerol minimal medium, starvation induced by glycerol depletion. (C) 8 μg/ml Polymyxin B and 4 μg/ml Colistin, added daily to cultures previously grown on glucose minimal medium (reference condition) with the first dose at 24h. Antibiotic concentration was 10x minimum inhibition concentration during growth (MIC). Each data point is the average of two biological repeats. Error bars are not shown because the variation is smaller than symbol size. Viabilities indicated at <10^1^ mean no colonies were detected from 100 μl culture. (D) Death rates for different Mg^2+^ concentrations. Reference condition (black circle in Fig. 1A) has 0.41 mM Mg^2+^. DNP+ cultures were supplemented with 1 mM 2,4-dinitrophenol during starvation in reference medium. Each data point is from a single experiment. All cultures were grown in glucose minimal medium. P-values (Student’s t-test, two-tailed) are indicated in comparison to the reference condition.

The physiological role of Lpp (Survival score: 7.2), together with OmpA (5.8), is to physically link outer membrane and cell wall. These proteins are essential for the cell envelope’s mechanical integrity and make up around half the cell envelope proteome and up to 12% of the total proteome (Fig. 2A). Motivated by this finding, we tested more cell envelope proteins, independent of their Survival scores, and found that starvation-survival is highly sensitive to knock-outs across different functionalities of the cell envelope (Fig. 2B), from outer membrane protein folding (BamE & Skp), cell wall synthesis and modification (Prc, NlpI, DolP & NlpD), phospholipid recycling (MlaA & MlaC) to cell envelope regulators (CpxA & RseA).

Conversely, only knockouts of porins, which facilitate diffusion of biomolecules across the outer membrane, have high Survival scores, and are highly abundant (Fig. 2A), did not change death rate (Fig. 2B).

This sheer abundance of cell envelope proteome makes it a compelling candidate for determining death rate in carbon starvation, as it would mean adaptation to optimal starvation survival would require a major reorganization of the proteome, rather than simply synthesizing a small set of proteins that could be achieved with a limited amount of stored nutrients. This could explain why the prior growth condition has such a big influence on starvation survival.

Many of the knock-outs that we identified are known to impair the mechanical integrity of the cell envelope (Bryant et al., 2020; Malinverni and Silhavy, 2009; McBroom et al., 2006; Rojas et al., 2018; Schäfer et al., 1999; Suzuki et al., 1978; Tsang et al., 2017; Yem and Wu, 1978). Therefore, we hypothesized that the mechanical integrity of the cell envelope plays a role in survival. To test this hypothesis, we applied antibiotics that are known to compromise the cell envelope, polymyxin B and colistin. Both antibiotics were highly effective when added 24 h into starvation, killing the entire population within 24 h and 48h, respectively (Fig. 2C). This is unusual, as most other antibiotics are ineffective on starving bacteria (Nguyen et al., 2011; Schink et al., 2022). Polymyxin B and colistin insert into the outer membrane between lipid A molecules and displace Mg^2+^ ions that shield repulsive negative charges between lipid A (Moore et al., 1986). Similarly, supplementing the starved culture with DNP, which acts as an ionophore and permeabilizes bacteria (McLaughlin, 1972) increased death rate 2.8-fold (Fig. 2D).

These results show that cell envelope integrity is essential for survival. In addition, we wanted to know if cell envelope integrity is also limiting for survival and tested the reverse effect by increasing Mg^2+^ concentrations. Indeed, we found a 40% drop in death rate at high Mg^2+^ concentrations compared to the reference condition (Fig. 2D), suggesting that cell envelope integrity is both essential and limiting.

### Ion homeostasis and cell envelope integrity account for the variation of starvation survival

But how does cell envelope integrity influence survival? We recently showed that the cost of ion homeostasis is a major determinant of lifespan of bacteria in carbon starvation (Schink et al., 2022). Based on this work, we hypothesize that changes in death rates in the CARLOS conditions were due to changes in the cost of ion homeostasis, mediated by the envelope proteome affecting membrane permeability. If membranes are less permeable, less energy is required to actively pump ions across the cytoplasmic membrane to remain ion homeostasis. This reduction in maintenance should then result in a lower death rate (Schink et al., 2019).

To test this hypothesis we used an ‘osmo-balanced’ medium with low salt concentrations, designed to minimize the content of inorganic ions that can diffuse across membranes (Schink et al., 2022). This medium reduces the death around 4-fold in standard glucose minimal medium conditions due to a reduction in maintenance rate required for ion homeostasis (Schink et al., 2022). If changes in the membrane permeability were responsible for the modulation of death rate in Fig. 1, we predict that the osmo-balanced medium should remove the growth-death dependence. In a perfect scenario, in which loss of ion homeostasis is the only possibility to die we would predict death rate to decrease to zero for all conditions, but we already know from previous work (Schink et al., 2022) that death rate does not decrease to zero with this medium, presumably because at some point lifespan will be limited by other causes.

On the other hand, if the growth-death dependence is independent of permeability of ions, for example if the modulated proteins and processes are preventing some type of damage or are influencing nutrient recycling, then we would expect the growth-death dependence to remain in ‘osmo-balanced’ medium but be shifted to a lower death rate.

Because the osmo-balanced medium allows us to distinguish whether the growth-death relation is due to a modulation of the ability to deal with ions in the medium, we grew *E. coli* in the CARLOS conditions followed by washing and resuspension in ‘osmo-balanced’ medium. We found that the characteristic growth rate dependence of the CARLOS condition was indeed abolished and no significant correlation between growth rate and death rate remained (Fig. 3). This confirms that differences of the cell envelope integrity and membrane permeability underlie the variations in death rate seen in Fig. 1A.

**Figure 3.**
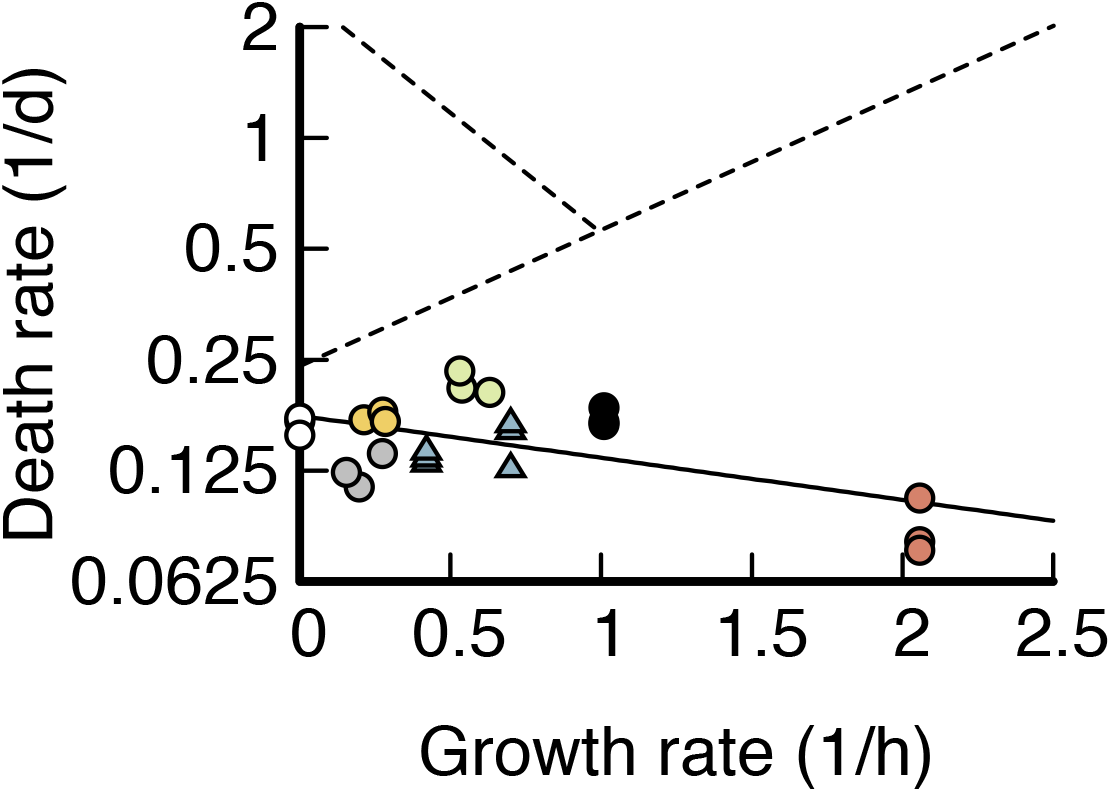
Death rates in osmo-balanced medium. Death rate plotted versus growth rate on a semi-log scale for experiments in ‘osmo-balanced’ medium. Colors indicate carbon limitation (C, ‘blue’, acetate and glycerol), anabolic limitation (A, ‘green’, 3μM IPTG), ribosome limitation (R, ‘yellow’, 4μM Chloramphenicol), rich medium (L, ‘red’, LB), LacZ overexpression (O, ‘grey’, 10ng/ml cTc), stationary phase (S, ‘white’) and the glucose reference condition (‘black’), as defined in Fig. 1A. The data shows weak anti-correlation with growth rate; solid black line is an exponential fit to the data (*R*^2^ = 0.34). Dashed lines are the exponential fits to the CARLOS data in Fig. 1A.

## Discussion

We performed an analysis of proteomics data to identify proteins, processes and cellular compartments that contribute to improved survival of carbon starvation. We identified cell envelope proteome and the membrane permeability as a key determinant of survival.

### Limitations of the study

Before discussing the implications of our work, we should spell out the limitations of our search for proteins that determine death rate. First, because we assume that the same proteins determine the modulation of death rate across all CARLOS conditions, we are prone to miss proteins that contribute to death rate in a single (or small subset) of conditions. Secondly, because we focused our analysis on the level of processes and cellular compartments, we will miss proteins and mechanisms that are not connected to significant processes and compartments. Therefore, these proteins could be undetected, even though they should in principle pop up with a high Survival-score (Table S2). It is thus possible that we missed additional mechanisms not connected to the cell envelope. Thirdly, the quality of the data sets varies across sources. The Schmidt et al data set, used for C, S and L, has a deep coverage that includes many low abundance proteins, and as a result, there is large number of proteins missing values in A and R (Table S2), which are derived from data sets with less coverage. This leads to some limitations in the scoring and the construction of the overall survival score. In order to retain the information available in the data set, we constructed our scores in such a way, that we omit missing values in the survival score, which leads to larger contributions of the available scores to the overall score. This can have two opposing types of effects: 1) Globally correlating proteins that are only measured in a subset of perturbations (which are generally of lower abundance) will have lower overall score, which introduces a bias towards proteins of high abundance. 2) If a protein has a strong anti-correlation in one of the missed conditions, the estimated survival score will be higher than it should be. To mitigate the effects of this in the downstream analysis, we perform our final enrichment analysis on the subset of proteins that were measured in at least 3 of the 5 CARLS conditions. In addition, proteins can be present in the data, but measured with so much underlying noise, that the direction of regulation cannot be determined with high confidence. This results in low Z-values and the effects are like the effects of missing values mentioned above. Finally, the number of individual data sets for each perturbations varies. While we compensate the impact of the number of data sets on the final Survival score by weighing the scores appropriately, more data sets per perturbation will reduce noise and improve the scoring. This creates a bias towards perturbations with many data sets, such as C, at the expense of those with few, such as A and R. It should be noted that the challenges of missing values, bias towards higher abundant proteins and measurement noise are common in the analysis of proteomics data and we have strived to find a compromise between stringency and comprehensiveness in the construction of our score.

Nevertheless, we have presented a variety of results that allow us to be confident in our main conclusions. In the construction of our score, we have shown good correlation between identical types of perturbations on independent data sets (Fig. EV1). We see enrichment in a canonical set of processes (Fig. 1C) and an increase in death rate for knockout of key genes in the cell envelope (Fig. 2B). We additionally validated that both weakening and strengthening the cell envelope affect death rate (Fig. 2C) and established that the growth-death relation can be abolished in the absence of ions in the medium (Fig. 3), which is a strong indication that the mechanism identified by our analysis, i.e., the integrity of the cell envelope, is the major contribution to the growth-death dependence of Fig. 1.

### Cost of building a strong cell envelope

Building a strong cell envelope is a costly endeavor. A third of the dry weight of a typical cell consists of cell envelope biomass, half of which consists of protein, and the other half phospholipids, LPS and cell wall (Neidhardt, 1996). Given the substantial fraction of biomass devoted to the cell envelope, it is reasonable that bacteria reduce this biomass investment into the cell envelope in favorable growth conditions, and instead focus on maximizing their growth rate. However, because of the magnitude of the biomass investment into the cell envelope, bacteria cannot easily adapt to starvation using carbon stores after sudden nutrient depletion, which would explain why pre-growth conditions have such a big effect on starvation survival, and why optimal survival requires prolonged adaptation.

### Role of the outer membrane in cell envelope integrity and starvation survival

Surprisingly, many of the proteins that we found to affect survival rates in carbon starvation are related to the outer membrane and overall, our data demonstrates an important role of the outer membrane for starvation survival. While the outer membrane is known to be a permeability layer, e.g., to antibiotics, it is less clear how it could achieve a similar effect for inorganic ions, which would explain improved survival in starvation. Ion export pumps, such as NhaA, are thought to only span the cytoplasmic membrane, meaning that ion gradients across the outer membrane should equilibrate. In this case, the outer membrane should offer little protection from inorganic ions. Indeed, in ‘plasmolysis’ from osmotic shocks, the inner membrane contracts as a result of osmotic pressure, while the outer membrane largely maintains its shape (Rojas et al., 2018), indicating that osmolytes readily diffuse into the periplasm. There is, however, considerable biochemical and mechanical interplay between inner and outer membrane that could contribute to permeability changes. Both membranes share phospholipids, and the outer membrane is involved in recycling lipids that have accumulated on the outer leaflet of the outer membrane (Malinverni and Silhavy, 2009). Recently, the outer membrane has been found to mechanically stabilize the inner membrane, e.g., via the tol-pal system, which is essential for bacteria to reach plasmolysis in starvation (Rojas et al., 2018; Shi et al., 2021). Integrity of the cell envelope could also protect periplasmic proteins from damage. We therefore argue that a mechanically strong cell envelope allows Gram-negative bacteria to stabilize and protect its cytoplasmic membrane, allowing bacteria to reduce their maintenance cost and increase their lifespan.

## Materials and Methods

### Strains

All strains used in this study are derived from wild type *E. coli* K-12 strain NCM3722 (Soupene et al., 2003). Strains NQ381 (attB:Plac-xylR, Km-Pu-lacY) was used for ‘catabolic limitation’ and was described in Ref. (You et al., 2013). In NQ381 strain the expression of the lactose transporter lacY is induced using 3-Methoxybenzamide (3MBA). NQ393 (ΔgdhA+plac-gltBD, attB(phage):lacIQ-tetR::Sp, ΔlacY) was used for ‘anabolic limitation’ and is described in Ref. (Hui et al., 2015). Using isopropylthio-β-galactoside (IPTG), the expression of gltBD is induced, which encodes glutamate synthase. NQ1389 (Ptet-tetR on pZA31; Ptetstab-lacZ on pZE1) was used to overexpress LacZ using chlortetracycline (cTc) as inducer and is reported in Ref. (Basan et al., 2015a). All knockouts shown in Fig. 2 were transferred from the Keio collection (Baba et al., 2006) to NCM3722 via P1 transduction to yield strain. Knockouts from Table S5 are taken from the Keio collection (Baba et al., 2006) and compared to the ancestor wild-type BW25113.

### Culture medium

The culture medium “N^−^C^−^ minimal medium” (Csonka et al., 1994), contains 1g K2SO4, 17.7 g K2HPO4, 4.7 g KH2PO4, 0.1 g MgSO4·7H2O and 2.5 g NaCl per liter. The medium was supplemented with 20 mM NH4Cl, as nitrogen source, and varying carbon sources. The ‘reference glucose condition’ contained 0.2% glucose. All chemicals were purchased from Sigma Aldrich, St. Louis, Mo, USA. The ‘osmo-balanced’ medium is described in (Schink et al., 2019), and contains 0.2 M MOPS (3-(N-morpholino)propanesulfonic acid), titrated to pH 7 with KOH, and 1 mM MgCl_2_, 0.1 mM CaCl_2_, 0.16 mM K_2_SO_4_, 0.5 mM K_2_HPO_4_, 22mM NH_4_Cl and lacks a carbon source. Cultures starved in the ‘osmo-balanced’ medium were previously grown in N-C-medium supplemented with NH_4_Cl and glucose.

### Culture conditions

Prior to each experiment, bacteria were streaked out from -80ºC glycerol stock on an LB agar plate supplemented with antibiotics if necessary. Bacteria were cultured in three steps. First, a seed culture was grown in lysogenic broth (LB) from a single colony. Second, the seed culture was diluted in N^−^C^−^ minimal medium supplemented with 20 mM NH_4_Cl and a carbon source and grown overnight for at least 5 doublings to exponential phase. The next morning, the overnight culture was diluted into fresh, pre-warmed N^−^C^−^ minimal medium supplemented with 20 mM NH_4_Cl and a carbon source and grown for another 5 to 10 doublings. At an optical density of 0.5 or below, the culture was washed by centrifugation (3 min at 3000 g) and resuspension into fresh, carbon-free, pre-warmed N^−^C^−^ minimal medium supplemented with 20 mM NH_4_Cl. This washing step removes excreted byproducts such as Acetate. For growth conditions known to fully respire carbon, e.g. wild-type NCM3722 grown on glycerol (Basan et al., 2015b), this washing step was omitted.

NQ381 was grown overnight and during the experimental culture with the inducer concentrations indicated in Table S1. NQ1389 was grown without inducer overnight, with indicated inducers added after dilution of the overnight culture, to prevent escape mutations and plasmid loss. NQ399 was grown with 100 μM IPTG overnight and the indicated IPTG concentrations after dilution of the overnight culture.

For culturing we used 20 mm x 150 mm glass test tubes (Fisher Scientific, Hampton, NH, USA) with disposable, polypropylene Kim-Kap closures (Kimble Chase, Vineland, NJ, USA) filled with 5 to 7 ml of medium. Cultures were sealed with Parafilm “M” (Bemis Company, Neenah, WI, USA) to prevent evaporation.

### Death rate measurements

Death rates were extracted using a linear fit of log-transformed viability measurements during the first 10 days of starvation, or until the viability reached below 10^7^ CFU/ml, whichever came first. Note that after the initial, exponential phase of starvation that we study, mutants take over, viability stabilizes, and cultures enter ‘long-term stationary phase’. For viability measurements, cultures were diluted in untreated, sterile 96 well plates (Celltreat, Pepperell, MA, USA) in three to four steps using a multichannel pipette (Sartorius, Göttingen, Germany) to a target cell density of about 4000 CFU/ml. 100 μl of the diluted culture was spread on LB agar plates supplemented with 25 μg/ml of 2,3,5-triphenyltetrazolium chloride to stain colonies bright red using Rattler Plating Beads (Zymo Research, Irvine, CA, USA), and incubated for 12 to 24 hours. Images of agar plates were taken with a Canon EOS Rebel T3i (Tokyo, Japan) mounted over an LED light box ‘Lightpad A920’ (Artograph, Delano, MN, USA), and analyzed using a custom script in Cell Profiler (Carpenter et al., 2006). Colony forming units per volume (CFU/ml) were calculated by multiplying the number of colonies per agar plate by the dilution factor.

### Stress conditioning

For pre-stressing, wild-type *E. coli* NCM3722 was grown in glucose minimal medium, either in a water bath at 40ºC (‘heat stress’), in medium supplemented with 50 mM NaCl (‘osmotic stress’) or in N^−^C^−^ medium adjusted to pH 6 using KOH (‘pH stress’). At an optical density OD6_00_ of about 0.5, cultures were washed and transferred to pre-warmed, carbon-free N^−^C^−^ supplemented with 20 mM NH_4_Cl, and the decay of viability was recorded for about 10 days.

### Anaerobic culturing

For anaerobic growth and starvation, cultures were grown in 0.05% glucose minimal medium in a vinyl anaerobic chamber (COY Lab Products, Grass Lake, Mi, USA), in Erlenmeyer flasks (Chemglass, Vineland, NJ, USA) on a magnetic stirrer (IKA RO10, Staufen, Germany), and not washed after the end of growth. For aerobic growth and anaerobic starvation, cultures were grown in 0.05% glucose minimal medium in an air incubator. At an optical density of about 0.5, cultures were centrifuged, supernatants were discarded, and pellets were introduced to the anaerobic chamber. In the anaerobic chamber, pellets were resuspended in pre-warmed, carbon-free minimal medium. All media were degassed prior to being introduced to the anaerobic chamber and left with open lid to be equilibrated for one week.

### Proteomics data processing

The MS proteomics data was downloaded from the corresponding PRIDE partner repositories (Schmidt et al.: PXD000498, Hui et al.: PXD001467, Houser et al.: PXD002140). For the Schmidt and Houser data sets, the .raw files were downloaded, for the Hui data set, peptide-intensity mappings were downloaded. The Schmidt and Hui data sets were searched MaxQuant (Cox and Mann, 2008) v. 1.5.7.4 using standard settings and “Label Free Quantification” (LFQ) and “Match between runs”. The Schmidt and the Houser data sets were searched against the reviewed Uniprot *E. coli* database (03/2019).

### Differential expression analysis of proteomics data

All data sets were processed with the MS-EmpiRe (Ammar et al., 2019) algorithm for differential quantification, which assigned fold changes and significance scores to each protein in a condition pair. For the Schmidt and Houser data sets, the MaxQuant “peptides.txt” and “proteinGroups.txt” files were used as input. The Hui data set was pre-processed as follows: As in the experimental setup of Hui et al. the data was quantified relative to a ^15^N-labelled spike-in, a direct assessment of LFQ values was not optimal. Similar to the approach of Ref. (Geiger et al., 2010), we first assessed the fold changes of each peptide relative to its heavy labeled spike-in. To preserve the intensity information, we re-scaled each spike-in fold change by the median intensity of the spike-in peptide over all conditions. This resulted in pseudo-intensities, which we further processed in a “LFQ-like” manner using MS-EmpiRe.

### Z-value based ranking of the proteome perturbations

As displayed in Fig. 1 and discussed in the main text, each growth condition has a corresponding death rate. Additionally, when we compare two conditions with one condition having a lower death rate than the other, we expect the lower death rate to be caused by increased expression of proteins. In our ranking, we hence had to identify proteins that show increased expression for decreased death rate over all conditions.

Our initial data set consisted of “Transporter Titration” (C), “Ribosome Limitation” (R) and “Anabolic Limitation” (A) in the Hui set, “Carbon Substrates” (C), “Chemostat” (C), “Rich Media” (L), “Stationary Phase” (S), “Osmotic Shock”, “Heat Stress” and “PH Stress” in the Schmidt set and “Stationary Phase” (S) in the Houser set. Each data set had a glucose reference condition.

The “Stationary Phase”, “Osmotic Shock”, “Heat Stress” and “PH Stress” perturbations were compared to the corresponding glucose reference using differential expression analysis with MS-EmpiRe. This assigned to each protein a log fold change denoting the change in expression and a significance score denoting the confidence in an expression change. The remaining perturbations were compared differently, because they consisted of multiple experiments with varying perturbation strength, e.g. different levels of transporter titration in the “Transporter Titration” data set. The individual conditions within a data set were ranked by their corresponding death rate and all combinations were compared in an increasing manner (e.g. c1 vs c2, c2 vs c3, c1 vs c3). This way, positive fold changes always correlate with better survival.

Concerning the Survival score calculation, our data was structured as follows: We have the five different CARLS perturbations, which we denote here as the set of perturbations *𝒫*. Each of the perturbations in *𝒫* has one or more data sets 𝒟, which are measurements of this perturbation type. For example, there are three data sets that belong to the perturbation of type C: different carbon sources (Data ref: Schmidt et al., 2016), the chemostat data set (Data ref: Schmidt et al., 2016), and the data set, where carbon uptake was regulated by transporter titration (Data ref: Hui et al., 2015). In other words, 𝒟_*C*_= {*𝒞*_carbon source,_ *𝒞*_chemostat_, *𝒞*_titration_}, where *𝒞* is the respective set of condition pairs. A condition pair (*α, β*) is the comparison of a condition *α* (e.g. glucose) against another condition *β* (e.g. acetate).

We started the calculation of the score at the level of a single condition pair and the score is calculated separately for each individual protein X. First, Box 1, step 1, we applied a Z-value transformation to a each condition pair (*α, β*) of protein X as follows:

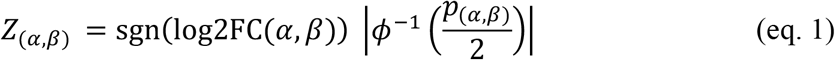

where log2FC is the log2 transformed fold change, p the p-value determined by MS-EmpiRe, sgn is the sign function and *ϕ*^−1^ is the inverse cumulative standard normal distribution function. The Z-value *Z*_(*α,β*)_ is the distance from zero in a standard normal distribution and can immediately be transformed back into a significance score. The higher the absolute value of the Z-value is, the more significant a measurement is. The Z-value carries both the direction of a regulation (via its sign) and its significance (via its value).

Next, Box 1, step 2, we merged the set of all condition pairs *𝒞* which belonged to the same data set D via the formula:

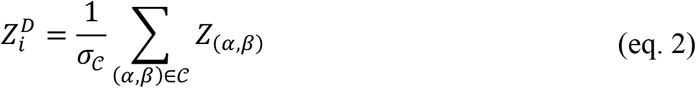

where 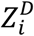 corresponds to the normalized Z value of the i-th data set. Normalization was performed by dividing by the standard deviation of the summed random variables denoted as *σ*_*𝒞*_, which is the square root of the variance.

Generally, the variance of the sum of multiple random variables is the sum over the corresponding covariance matrix. As we combine variables that follow a standard normal distribution, we can derive the individual covariances in our setup as follows: When random variables from two independent standard normal distributions are compared, (e.g. Z1 comes from the comparison C1 vs. C2 and Z2 comes from the comparison C3 vs. C4) the covariance is equal to 0. When random variables that share one sample are compared (e.g. Z1 comes from the comparison C1 vs. C2 and Z2 comes from the comparison C1 vs. C3) it can be shown that the covariance is equal to 0.5 (Ammar et al., 2019). This then allows us to determine *σ*_*𝒞*_ and therefore 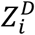.

Subsequently, Box1, step 3, we merged the datasets 𝒟 that were available for each type of perturbation *P* (i.e. each of the CARLS perturbations). Depending on the perturbation, the number of available datasets varied – for example, for the C perturbation, we had three datasets (𝒟_*C*_ = {*C*_carbon sources_, *C*_chemostat_, *C*_titration_}), while for the A perturbation, we had only one dataset (𝒟_*A*_ = {*A*_titration_}). To obtain a normalized Z value for each perturbation, we used the equation

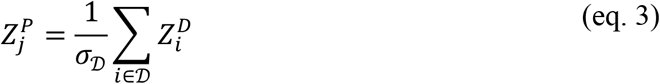

where 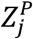 corresponds to the normalized Z-value of j-th Perturbation P. The value is normalized by the standard deviation over the datasets 𝒟, denoted as *σ*_𝒟_. The calculation is again carried out via the summation over the covariance matrix. However, because the random variables 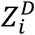 are independent from each other, all covariances are zero except for the covariances of the random variables with themselves (i.e. their variances). Because each 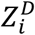 is (0,1) normalized according to eq. 2, the resulting variance corresponds simply to the number of elements in 𝒟 and *σ*_𝒟_ is therefore the square root of the number of elements *n* in 𝒟. In the case that 𝒟 contains only one element, for example in the 𝒟_*A*_ case, 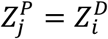.

For the final calculation of the Survival score Box1, step 4, the set of perturbations *𝒫* is summarized to the Survival score *Z*^*S*^ via the equation:

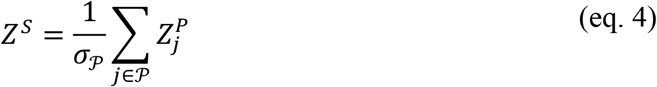

The standard deviation *σ*_*𝒫*_ is the square root of the number of elements in *𝒫*, following the same reasoning as above. The overall score is robust against missing values, as perturbations that are not available (e.g. if for a certain protein X, we have only *𝒫* = {C, R, L, S} available) also do not contribute to *σ*_*𝒫*_. The available Z-values then contribute more strongly to the overall score.

### Absolute quantification of proteins

For absolute quantification, we used protein synthesis rates derived by (Li et al., 2014) from ribosomal sequencing data of a MG1655 glucose reference condition. Synthesis rates were used as proxies for copy numbers and multiplied by the respective molecular weights to obtain mass estimates. Further conditions were compared relatively to the reference with MS-EmpiRe and the mass estimates were scaled by the respective fold changes. To determine the mass fraction of a gene set, the genes of the set were summed and divided by summed mass of all genes.

### GO enrichment analyses

The Gene Ontology (GO) was downloaded from http://geneontology.org (03/2019) together with the *E. coli* “ecocyc.gaf” annotation. The relations “is_a” and “part_of” were used for the construction of the gene sets. Score-based analysis was carried out using the Kolmogorov-Smirnov test with signed scores. Multiple testing correction was carried out via the Benjamini-Hochberg procedure.

## Acknowledgements

This project was supported by MIRA grant (5R35GM137895) and an HMS Junior Faculty Armenise grant to MB. SJS was supported by EMBO via a Long-term fellowship (ALTF 782-2017) and HFSP via a long-term fellowship (LT000597/2018). CA was supported by a PhD scholarship of the Deutsche Forschungsgemeinschaft (DFG, Graduate School QBM).

## Author contribution

SJS, CA and MB conceived the project. SJS and CA performed experiments and proteomics analysis. YFC constructed strains. RZ supervised proteomics analysis. SJS, CA, RZ and MB wrote the paper.

## Data availability

The uploaded Source Data contains the data underlying each figure (“Figure Data.zip”). The proteomics raw data can be found under the repositories specified in the references. Code and data for bioinformatics analyses are available on Github (https://github.com/ammarcsj/ecoli_survival_scoring).

## Expanded View Figures Legends

**Figure EV1.**
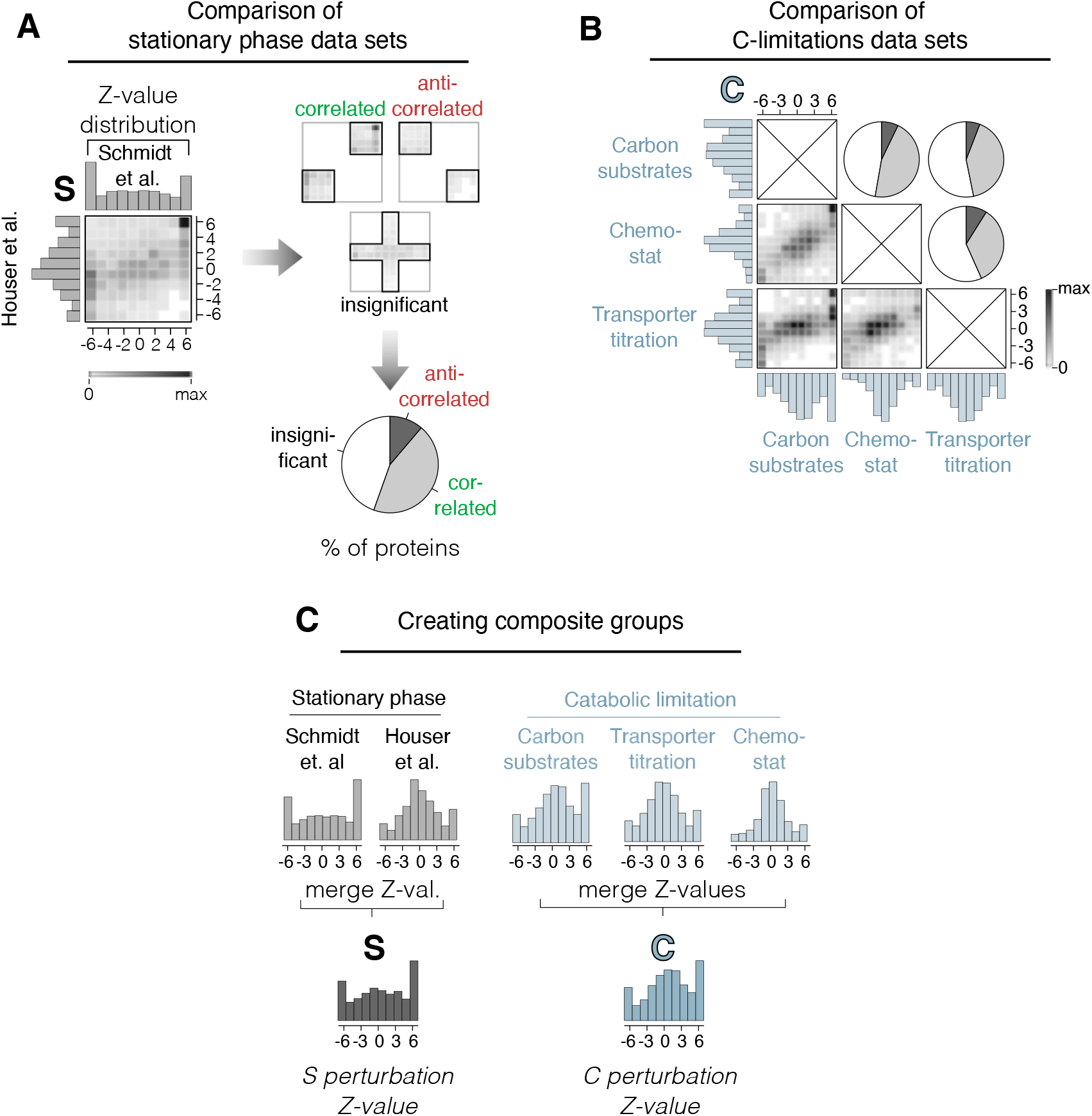
Comparison and merging of MS proteomics data from different sources. (**A**) Comparison of Z-value distributions from two different data sets, by Houser et al. (Data ref: Houser et al., 2015) and Schmidt et al. (Data ref: Schmidt et al., 2016), both measured in stationary phase after growth on glucose. On left: Z-value distributions of individual experiments are shown on left and top of correlogram. Shaded area inside the correlogram depicts frequency distribution of individual proteins that are measured in both data sets. On right: Proteins scored in top right and bottom left corner are counted ‘correlated’, proteins in the top left and bottom right are counted as ‘uncorrelated’. Cut-off for significance is chosen at Z = 1.28, corresponding to a p-value of 0.1. Proteins with at least one Z-value less than 1.28 are counted as uncorrelated. On bottom: Quantitative analysis of correlation. Only a minority of proteins are observed to anti-correlate between data sets, with the majority either correlating or being statistically insignificant. (**B**) Analysis of correlograms and quantification in pie charts of three data sets of different catabolic limitation analog to panel A. ‘Carbon substrates’ and ‘chemostat’ taken from Schmidt et al (Data ref: Schmidt et al., 2016), ‘Transporter titration’ taken from Hui et al (Data ref: Hui et al., 2015). Different types of catabolic limitation show high correlation between data sets. (**C**) Data sets from panel A and B, respectively, are merged to form a single Z-value distribution for each growth perturbation.

**Figure EV2.**
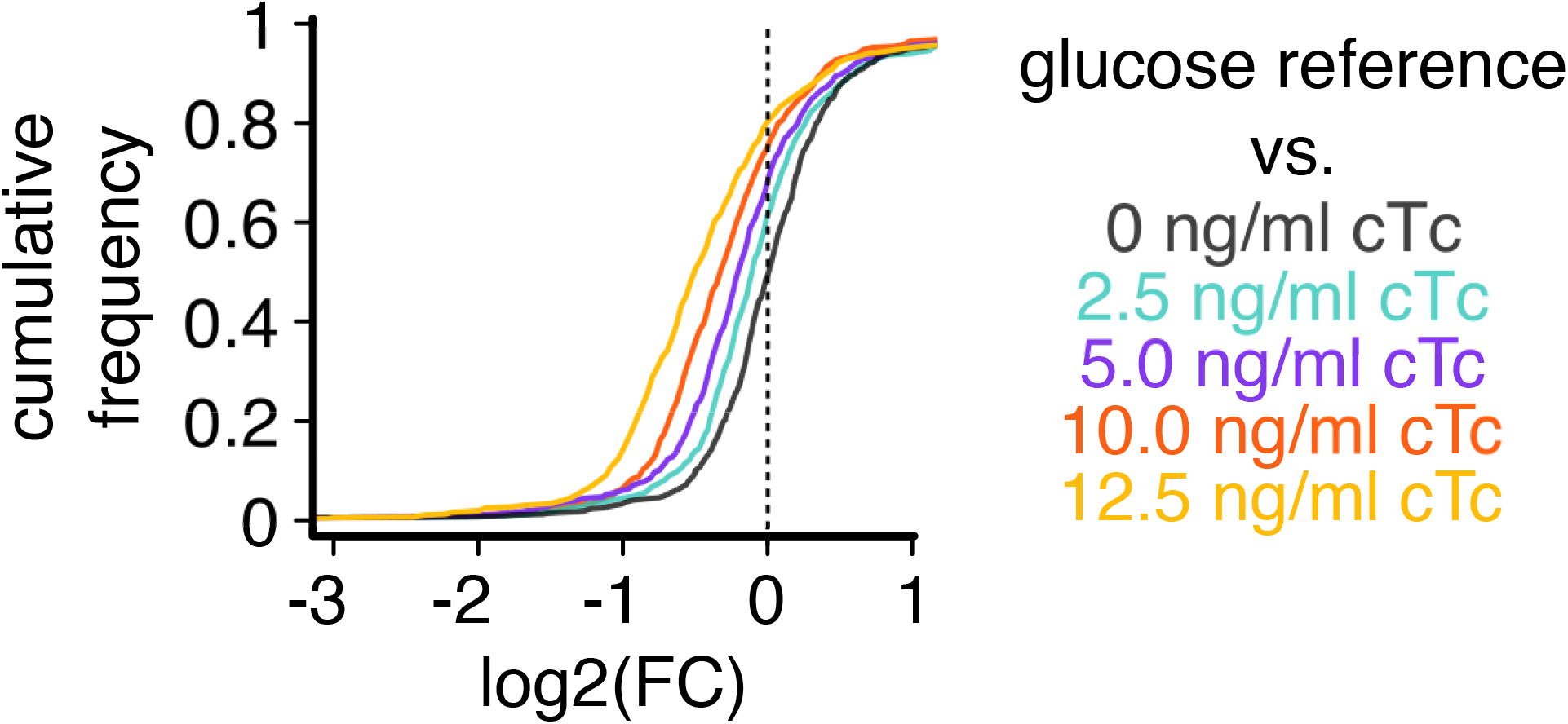
Cumulative distribution of fold-changes under expression of a useless protein (O condition). Fold-change of protein abundances (FC) in LacZ overexpression using strain NQ1389 compared to the reference condition of wild-type *E. coli* grown on glucose. The abundance of vast majority of proteins decreases with induction of LacZ (inducer: chlortetracycline, cTc).

**Figure EV3.**
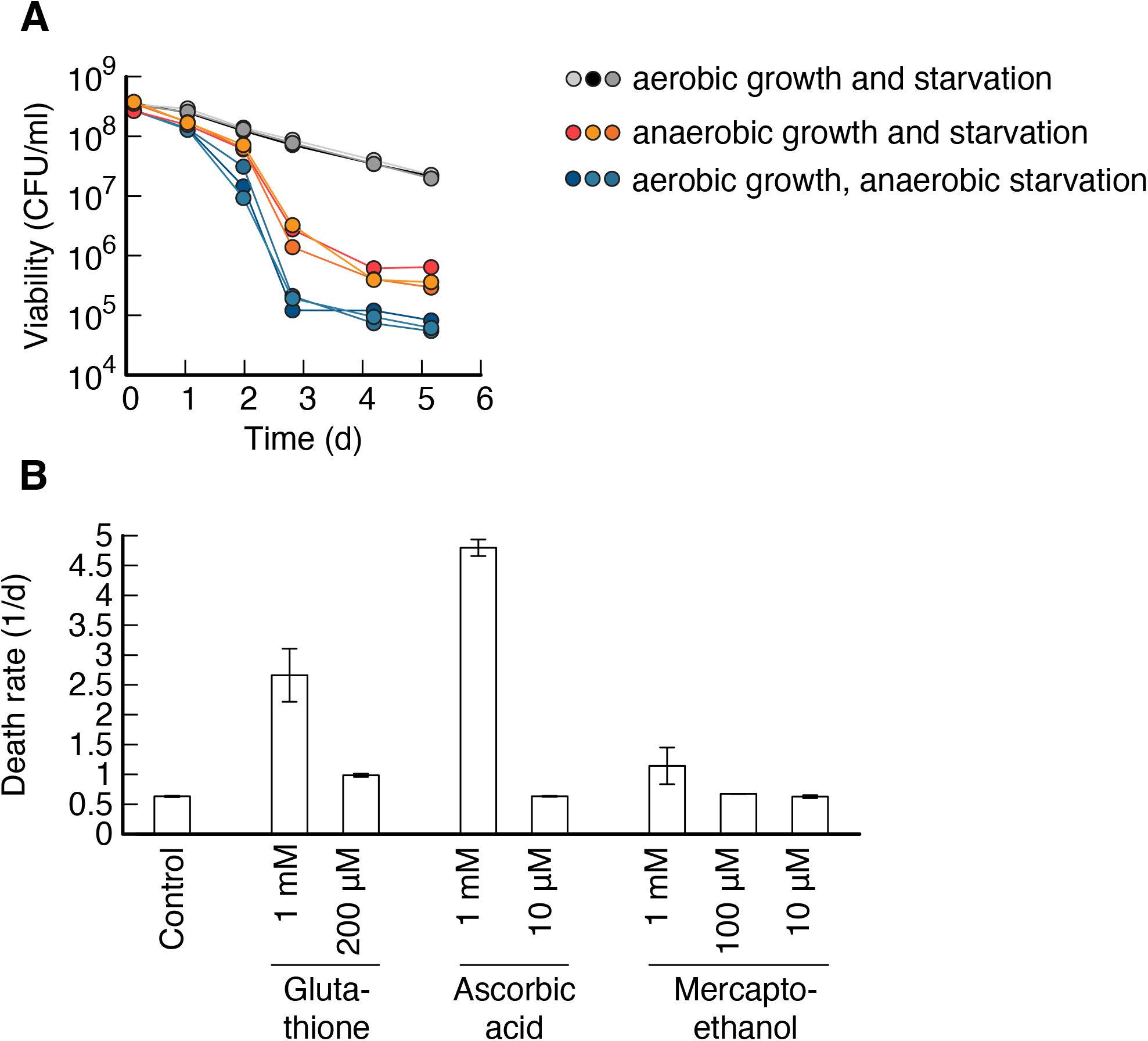
Starvation in the absence of oxidative damage. (**A**) Viability over time in E. coli cultures starved in either anaerobic or aerobic conditions. Cultures previously grown on glucose minimal medium. Neither performing both starvation and growth (orange), nor only starvation in anaerobic conditions (blue) decreased death rate. Instead in both cases, viability decreased several orders of magnitude more compared to aerobic conditions. This finding is in disagreement with previous claims that anaerobic starvation prevents loss of viability (Dukan and Nyström, 1999), but matches the observation that bacteria actively recycle nutrients from dead cells (Schink et al., 2019), which is more efficient if bacteria can use respiration. (**B**) Starvation in the presence of antioxidants glutathione, ascorbic acid (Vitamin C) and beta-mercaptothanol. High concentrations of antioxidants increase death rate, while low concentrations show no significant effect over control. No antioxidant condition significantly decreased death rate.

**Figure EV4.**
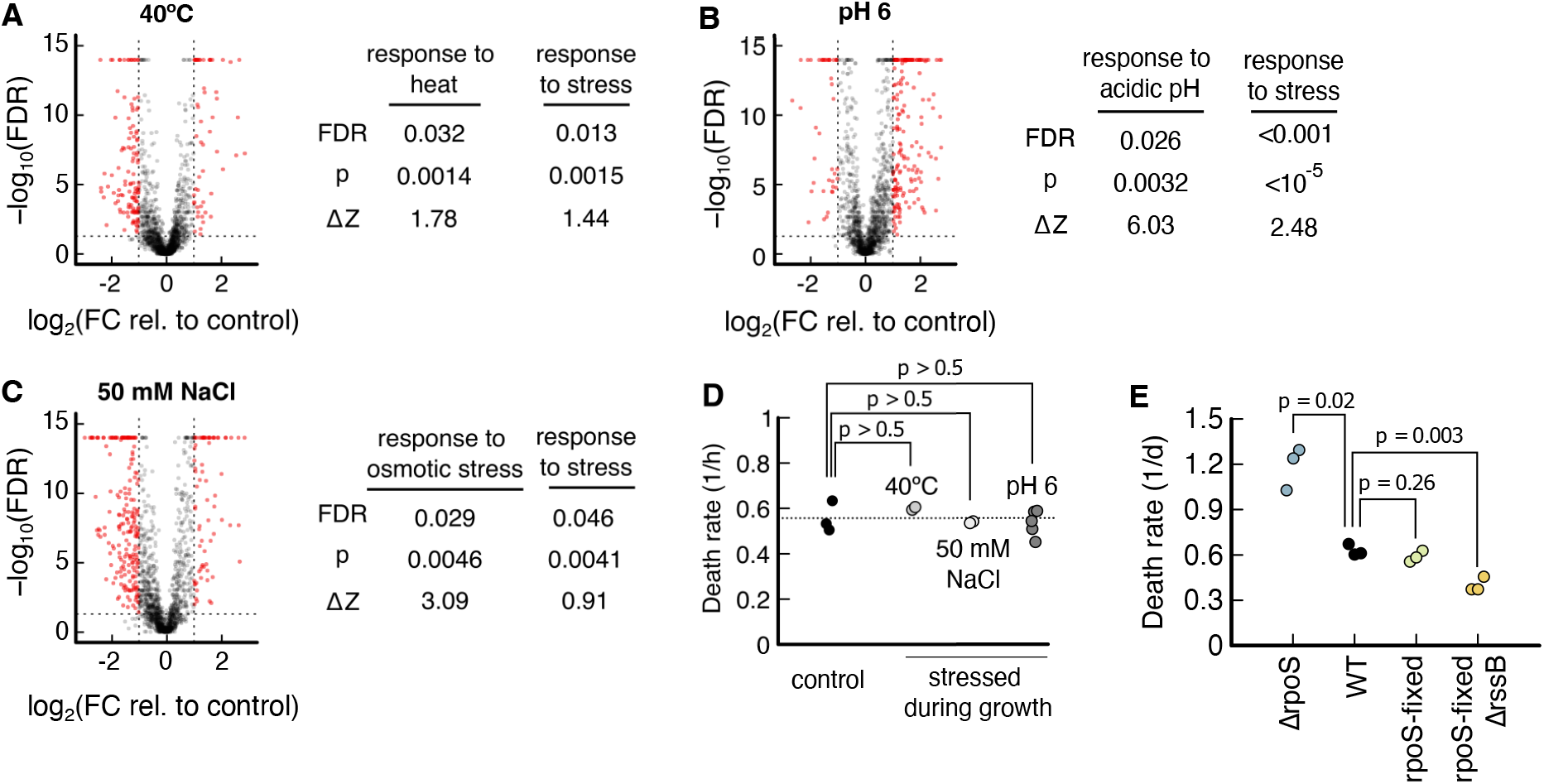
Effect of pre-stressing on proteome and survival kinetics. (**A**-**C**) Comparison of stress conditions to a glucose reference. On left side of each panel, volcano plots of individual proteins, showing the probability of being a response versus the logarithm of the fold change. Proteins with fold change higher than 2, i.e., *log*_2_(2) = 1, and false discovery rate smaller than 0.05, i.e. *log*_10_(0.05) = −1.3, are colored in red. On the right side of the panel, for each pre-stress (40ºC, pH 6 and 50 mM NaCl), the corresponding stress response and the general ‘response to stress’ are tested for significant upregulation using Kolmogorov-Smirnov tests. In each pre-stress, both the specific and the general stress response are significantly upregulated, FDR < 0.05. Data source: (Data ref: Schmidt et al., 2016). (**D**) After stressing during growth, bacteria are transferred to pre-warmed, carbon-free minimal medium without stress. Death rates of neither pre-stressing condition led to a significant change in death rate, p> 0.5. (**E**) Death rate of wild-type NCM3722 compared to ΔrpoS, a ‘rpoS-fixed’ strain where an amber stop codon in *rpoS* is restored (Mori et al., 2021) and a strain with ‘rpoS-fixed’ and ΔrssB. All mutants have NCM3722 as background. p-values in panels D and E are calculated using a two-tailed t-test.

**Figure EV5.**
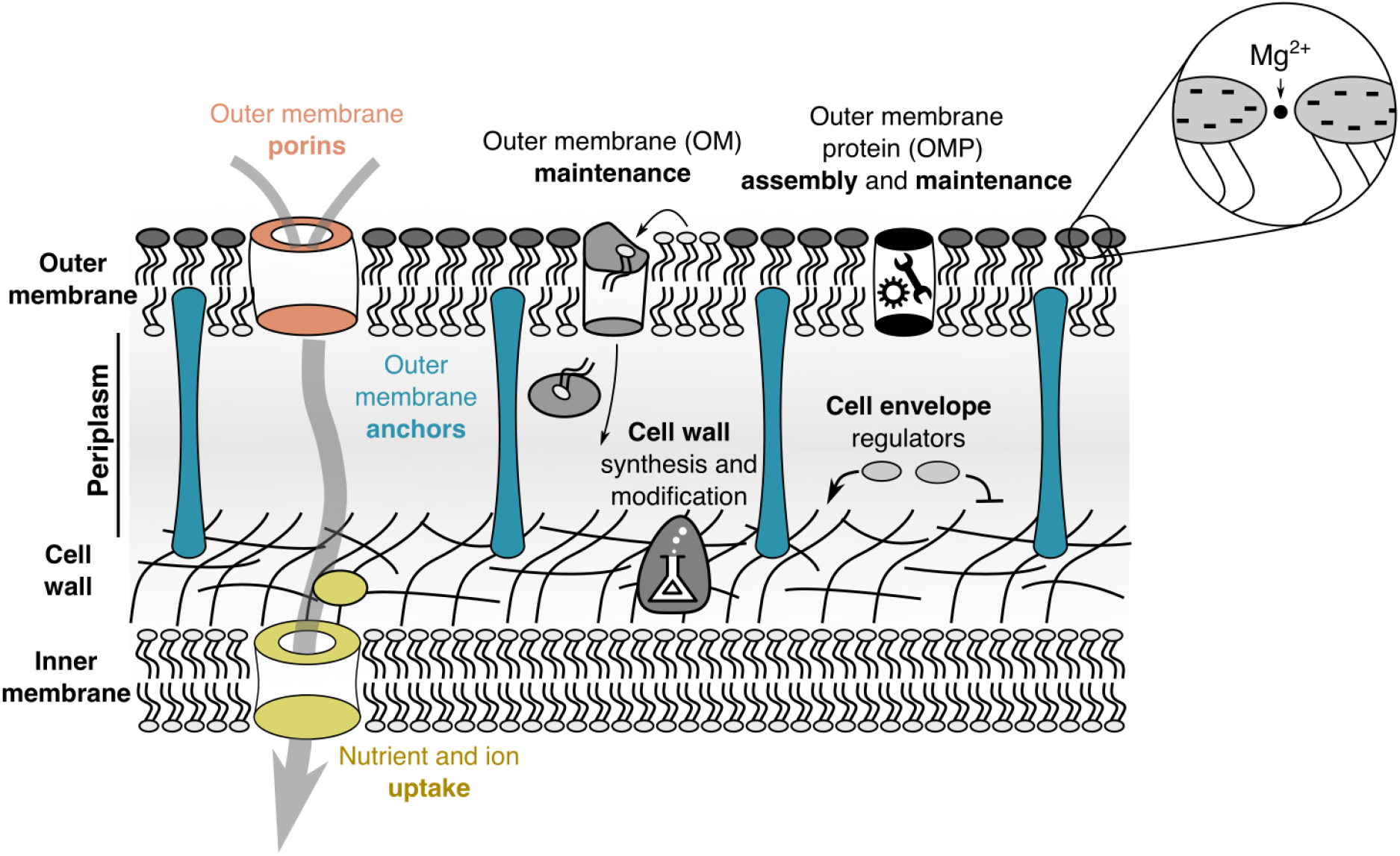
The cell envelope of *E. coli*. For gram-negative bacteria, the cell envelope consists of the outer membrane, the cell wall, and space between inner and outer membrane called periplasm. Proteins in the cell envelope have diverse functions. Outer membrane anchors (blue), specifically Lpp and OmpA, connect the outer membrane to the cell wall. Outer membrane porins (red) are large tunnel-shaped proteins that facilitate diffusion of biomolecules and ions across the outer membrane. Nutrient uptake systems (yellow) import biomolecules and ions across the inner membrane. While most of the nutrient uptake system is in the inner membrane, thus not considered as part of the envelope, binding proteins (yellow circle in periplasm) are. There are over 100 different kinds of binding proteins in *E. coli*, which make up a significant fraction of the proteome of the cell envelope. Other proteins include proteins involved in maintenance of the outer membrane, e.g. the Mla system which shuffles phospholipids from the outer leaflet of the outer membrane to the inner membrane. Outer membrane protein (OMP) assembly and maintenance are proteins that fold outer membrane proteins, e.g. Bam complex, proteases which degrade misfolded proteins, e.g. DegP, and chaperones that prevent folding of outer membrane proteins in solution, e.g. Skp. Cell envelope regulators include the CpxAR signal transduction system and regulation of RpoE via anti-sigma factor RseA. Cell wall synthesis and modification includes cell wall hydrolases and regulators of cell wall synthesis. Lipid A, the lipid on the outer leaflet of the outer membrane is highly charged and requires Mg2+ ions for shielding (Schneck et al., 2010).

## Supporting Information Tables

All Supporting Information Tables are provided in the single excel file “SupportingInformationTables.xlsx” with individual sheets for Tables S1 to S5.

**Table S1**. Growth and death rates in different growth perturbations. Coloring corresponds to Fig. 1. For all conditions listed here, cultures were washed and resuspended in carbon-free medium. All strains are derived from wild-type *E. coli* K-12 NCM3722. The titratable glutamate synthesis strain NQ393 contains a deletion of gdhA and the promotor of gltBD is replaced by P_lac_, which is induced with Isopropyl β-D-1-thiogalactopyranoside (IPTG) (Hui et al., 2015). Strain NQ1389 contains two plasmids, P_tet_-tetR on pZA31 and P_tetstab_-lacZ on pZE1, induced by Chlortetracycline (cTc), allowing high levels of superfluous LacZ expression (Basan et al., 2015a). In addition to Fig. 1A, the table also includes data from a titratable lactose uptake strain NQ381 has the P_lac_ replaced by Pu, which is induced with 3-methylbenzyl alcohol (3MBA) (You et al., 2013). This data follows the same trend as other carbon limitations but is not shown in Fig. 1A.

**Table S2**. List of Z values, Survival scores and fold-changes of abundance in pre-stress conditions. A total of 2039 proteins were scored. Values indicated with NA are not available. See Methods for details on the calculation of Z values and Survival scores. The CARLS columns show the Z values derived as described in the methods. For the C and the S conditions, several data sets were available, which were merged as visualized in Figure S2. The Z values for the individual data sets are also listed as follows: C_chemostat is the chemostat conditions of (Data ref. Schmidt et al., 2016), C_carbon_sources is the different carbon growth conditions of (Data ref: Schmidt et al., 2016), C_glucose_uptake_titration is the glucose transporter titration set from (Data ref: Hui et al., 2015) ‘S_houser’ is the Z values after 48 h of stationary phase respectively in the study of (Data ref: Houser et al., 2015) ‘S_schmidt’ is the Z values of the stationary phase condition of. (Data ref: Schmidt et al., 2016). ‘HEAT’, ‘OSM’ and ‘pH’ are Z values of pre-stress conditions with 42ºC, 50mM NaCl and pH 6, respectively, of (Data ref: Schmidt et al., 2016). ‘FC CHvsOSM’, ‘FC CHvsHEAT’ and ‘FC CHvsPH’ are the fold-changes of protein abundances in pre-stress conditions compared to reference glucose conditions, which are used in Fig. EV4.

**Table S3**. Full list of GO processes and GO compartments and their False discovery rates (FDR) and median Survival scores.

**Table S4**. Absolute abundances of cell envelope proteins, and protein groups in stationary phase, glucose reference condition and LB. See Methods for details on the quantification.

**Table S5**. Results from a knock-out screen. Genes were chosen to have high Survival scores and to be non-essential. Note that these genes were selected based on a preliminary version of the proteomics analysis, meaning that they do not 100% overlap with the highest scoring set of genes of Table S2. Survival scores of the final analysis are given in column D. Cultures were grown in N-C-supplemented with NH_4_Cl and 20mM glycerol in 96 well plates and starved by letting glycerol deplete. Death rate is estimated from a measurement of viability after glycerol ran out and a measurement after five days of starvation. Each data point is from a single experiment, and the hits were independently verified (Fig. 2). Out of the top five genes, three (*lpp, bamE* and *mlaC*) are essential for the cell envelope integrity.

